# Altered neurological and neurobehavioral phenotypes in a mouse model of the recurrent *KCNB1*-p.R306C voltage-sensor variant

**DOI:** 10.1101/2023.03.29.534736

**Authors:** Seok Kyu Kang, Nicole A. Hawkins, Dennis M. Echevarria-Cooper, Erin M. Baker, Conor J. Dixon, Nathan Speakes, Jennifer A. Kearney

## Abstract

Pathogenic variants in *KCNB1* are associated with a neurodevelopmental disorder spectrum that includes global developmental delays, cognitive impairment, abnormal electroencephalogram (EEG) patterns, and epilepsy with variable age of onset and severity. Additionally, there are prominent behavioral disturbances, including hyperactivity, aggression, and features of autism spectrum disorder. The most frequently identified recurrent variant is *KCNB1*-p.R306C, a missense variant located within the S4 voltage-sensing transmembrane domain. Individuals with the R306C variant exhibit mild to severe developmental delays, behavioral disorders, and a diverse spectrum of seizures. Previous *in vitro* characterization of R306C described loss of voltage sensitivity and cooperativity of the sensor and inhibition of repetitive firing. Existing *Kcnb1* mouse models include dominant negative missense variants, as well as knockout and frameshifts alleles. While all models recapitulate key features of *KCNB1* encephalopathy, mice with dominant negative alleles were more severely affected. In contrast to existing loss-of-function and dominant-negative variants, *KCNB1*-p.R306C does not affect channel expression, but rather affects voltage-sensing. Thus, modeling R306C in mice provides a novel opportunity to explore impacts of a voltage-sensing mutation in *Kcnb1*. Using CRISPR/Cas9 genome editing, we generated the *Kcnb1^R306C^* mouse model and characterized the molecular and phenotypic effects. Heterozygous and homozygous R306C mice exhibited pronounced hyperactivity, altered susceptibility to flurothyl and kainic acid induced-seizures, and frequent, long runs of spike wave discharges on EEG. This novel model of channel dysfunction in *Kcnb1* provides an additional, valuable tool to study *KCNB1* encephalopathies. Furthermore, this allelic series of *Kcnb1* mouse models will provide a unique platform to evaluate targeted therapies.

## 1. Introduction

Disease phenotyping in animal models is an important step for understanding many aspects of human physiological and pathological processes. This is especially true for neurological diseases that are often rare and complex, because the manifestation of disease phenotypes involves advanced dimensions of human biology such as the collective environment (i.e. neural circuitry) and time (i.e. neurodevelopment), which cannot be adequately addressed in non-animal experimental systems like cell culture or computational methods (Chesselet and Carmichael, 2012). CRISPR/Cas9 genome editing technology has enabled faster and more accurate recapitulation of human diseases in mice, and thus accelerated construction of genotype-phenotype correlations for rare diseases with genetic basis (Aida et al., 2014; Platt et al., 2014).

*KCNB1* encephalopathy is a rare autosomal dominant disorder caused by pathogenic variants in the *KCNB1* gene that most often arise de novo in the affected child. Individuals with *KCNB1* encephalopathy present with global developmental delay in infancy or early childhood accompanied by features of autism spectrum disorder, abnormal EEG patterns, and development of epilepsy in most children, although epilepsy severity and treatment response are variable (Bar et al., 2020a; Bar et al., 2020b; de Kovel et al., 2017; Kang et al., 2019; Saitsu et al., 2015; Scheffer and Liao, 2020; Thiffault et al., 2015; Torkamani et al., 2014). *KCNB1,* encoding the KV2.1 voltage-gated potassium channel alpha subunit, is a critical contributor to neuronal repolarization and homeostasis (Murakoshi and Trimmer, 1999). The majority of *KCNB1* variants studied thus far in heterologous cells and cultured neurons have been shown to confer an ultimate outcome of loss-of-function (LoF) that prevents the channel from conducting K^+^ ions across the plasma membrane (Kang et al., 2019; Saitsu et al., 2015; Thiffault et al., 2015).

Although the ultimate outcome is largely LoF, there are several classes of underlying mechanisms that lead to diminished channel function, including defective KV2.1 synthesis, trafficking, or function. Targeted therapies tailored to the specific mechanism can improve patient outcomes (Haq et al., 2022; Tian et al., 2022), highlighting the important of representing these different mechanisms in preclinical animal models. Variants modeled in mice to date have focused on those that affect KV2.1 expression, including *Kcnb1^G379R^*, *Kcnb1^R312H^* and *Kcnb1* null and frameshift alleles (Bortolami et al., 2022; Hawkins et al., 2021; Speca et al., 2014). *KCNB1*-p.G379R exhibited a dominant-negative LoF phenotype with altered ion-selectivity in CHO-K1 cells (Torkamani et al., 2014), and recapitulated dominant-negative cellular phenotypes, epilepsy, background EEG abnormalities and neurobehavioral symptoms in mice (Hawkins et al., 2021). *KCNB1*-p.R312H exhibited a LoF phenotype with deficient cell surface expression in CHO-K1 cells, and diminished KV2.1 protein and seizures in knock-in mice (Bortolami et al., 2022; Kang et al., 2019).

*KCNB1*-p.R306C represents another major class of channel dysfunction, diminished function due to altered voltage-sensing, and is one of the most recurrent variants (Kang et al., 2019). At least six cases with *KCNB1*-p.R306C have been described in the literature and severity of the associated phenotypes ranges from mild developmental disability with absence epilepsy to severe developmental disability with intractable epilepsy that includes multiple seizure types (Bar et al., 2020a; de Kovel et al., 2017; Kang et al., 2019; Saitsu et al., 2015). Arginine 306 is one of the positively charged residues in the S4 transmembrane domain that is critical for voltage-sensor function of the KV2.1 channel (Catacuzzeno and Franciolini, 2022). Functional studies of the R306C variant in heterologous expression systems showed normal cell surface expression, but loss of KV2.1 channel current and shifts in voltage-dependence of activation (Kang et al., 2019; Saitsu et al., 2015). Consistent with this, transient overexpression in primary cultured cortical pyramidal neurons resulted in lower sensitivity and cooperativity of the voltage sensor and severely impaired capacity for repetitive firing (Saitsu et al., 2015). To model KV2.1 voltage-sensor dysfunction *in vivo*, we generated and characterized a novel *Kcnb1^R306C^* mouse line using CRISPR/Cas9 genome editing (Platt et al., 2014). We evaluated the effects of heterozygosity and homozygosity for the R306C variant on KV2.1 expression and localization in cultured neurons, as well as seizure susceptibility, EEG, and locomotor activity in behaving mice. Although expression and localization of KV2.1 was unaffected, *Kcnb1*^R306C^ mice displayed hyperactivity, altered susceptibility to induced seizures by flurothyl and kainic acid (KA), and significant EEG abnormalities. Thus, *Kcnb1^R306C^*mice recapitulate key features of *KCNB1* encephalopathy and will be a useful platform for studying disease mechanisms, probing variable expressivity, and evaluating potential therapies.

## 2. Methods

### 2.1 Mice

Gene editing was performed in fertilized C57BL/6J embryos via electroporation of CRISPR components. The Cas9 protein was complexed with a CRISPR guide RNA (gRNA), which generated DNA double strand break in exon 2 of the *Kcnb1* gene. A single-stranded donor oligonucleotide (ssODN) was included in the reaction to introduce the R306C point mutation via homology dependent repair (HDR): gRNA sequence: 5’ GGGCCAACTTCAGGATGCGC 3’ The ssODN sequence is as follows: 5’CTGCGCAGCGTGAAGCCCAAGGACTGCAGACCGGTGGAGTGGCGGGCCAACTTCAGGATGCaCAGaATGCGCATGATGCGGAAGATCTGGACCACACGGCGCACATTCTGGA ACTGCAGC 3’. This was designed as an antisense sequence, complementary to the sense (coding) strand. Highlighted in green is a change in the coding sequence from CGC→tGC, resulting in the *Kcnb1* R306C point mutation upon ssODN mediated HDR. Highlighted in teal is a silent mutation changing coding of the Ile from ATC-ATt which disrupts the protospacer adjacent motif (PAM,-NGG) site preventing re-cutting of the site once ssODN mediated HDR occurs.

For the CRISPR components we used the AltR-Cas9 system (Integrated DNA Technologies, Inc. (IDT), Coralville, Iowa). Briefly, the gRNA was custom synthesized as crRNA and complexed with tracrRNA (IDT, 1070532) to form the gRNA complex. It was combined with HiFidelity Cas9 protein (IDT, 1081064) to form the ribonucleotide protein complex (RNP). The 120 nucleotide ssODN (IDT) was added to the reaction mixture after RNP formation, prior to electroporation. The final concentration of the electroporation mixture was 3uM Cas9, 3uM gRNA, 10uM ssODN.

Potential founders were screened by PCR of genomic DNA with primers outside of the homology region for the repair oligo (Table 1). PCR products were cloned into pCR-TOPO (ThermoFisher) and Sanger sequenced. The mosaic R306C founder was backcrossed to C57BL/6J mice (Jackson Labs, #000664, Bar Harbor, ME) to generate N1 offspring. The N1 offspring were genotyped by Sanger sequencing to confirm transmission of the R306C editing event and absence of off-target events at predicted sites with <3 mismatches. N1 males with the confirmed on-target event and without predicted off-target events were bred with C57BL/6J females to establish the line *Kcnb1^em3Kea^* (MGI: 675522), which has been maintained as an isogenic strain on C57BL/6J by continual backcrossing of *Kcnb1^R306C^*heterozygous mice (abbreviated *Kcnb1^C/+^*) with inbred C57BL/6J mice. For experiments, male and female *Kcnb1^C/+^*mice were intercrossed to generate *Kcnb1^+/+^* wildtype (WT), heterozygous *Kcnb1^C/+^* (C/+), and homozygous *Kcnb1^C/C^* (C/C) mice.

**Table 1.**
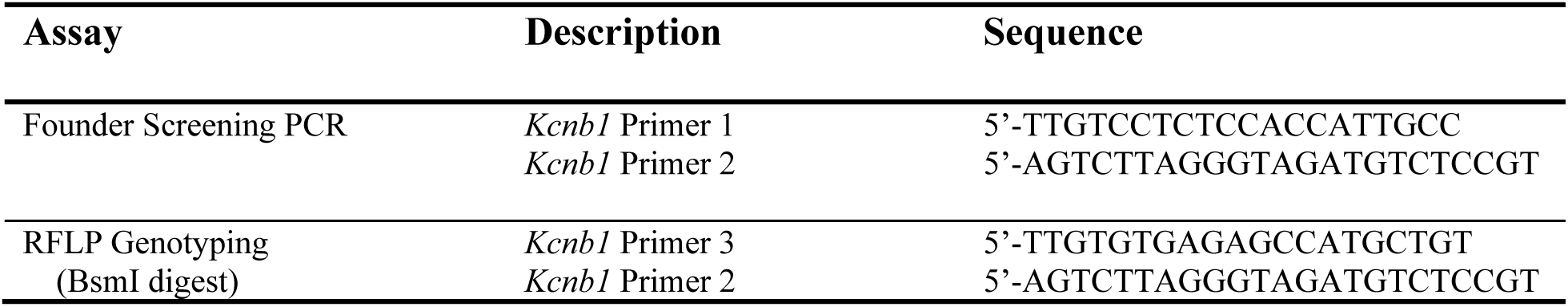
List of Primers

Mice were maintained in a Specific Pathogen Free (SPF) barrier facility with a 14-h light/10-h dark cycle and access to food and water *ad libitum*. Both female and male mice were used for all experiments. All animal care and experimental procedures were approved by the Northwestern University Animal Care and Use Committees in accordance with the National Institutes of Health Guide for the Care and Use of Laboratory Animals. The principles outlined in the ARRIVE (Animal Research: Reporting of in vivo Experiments) guideline was considered when planning experiments (Percie du Sert et al., 2020).

### 2.2 Genotyping

Mice were genotyped by PCR of genomic DNA isolated from tail biopsies, using a restriction fragment length polymorphism (RFLP) assay. Genomic DNA was amplified using RFLP genotyping primers (Table 1), followed by restriction digest with BsmI for at least 15 minutes at 65°C. Digestion resulted in 402 bp and 253 bp products for the mutant allele and 655 bp for the WT allele.

### 2.3 Transcript Analysis

Forebrain isolated from male and female WT, *Kcnb1^C/+^* and *Kcnb1*^C/C^ mice at postnatal days 58-83 (P58-83) was used for total RNA extraction using TRIZol reagent according to the manufacturer’s instructions (Invitrogen). First strand cDNA was synthesized using 4 μg of RNA using oligo(dt) primer and Superscript IV according to the manufacturer’s instructions (Life Technologies). First strand cDNA samples were diluted 1:10 and 5 μL was used as template with ddPCR Supermix for Probes (No dUTP) (Bio-Rad) and TaqMan Gene Expression Assays (Life Technologies) for mouse *Kcnb1* (FAM-MGB-Mm00492791_m1) and *Tbp* (normalization control; VIC-MGB-Mm00446971_m1). Reactions were partitioned into a QX200 droplet generator (Bio-Rad) and then amplified using PCR conditions: 95 °C for 10 min, 44 cycles of 95 °C for 30 s and 60 °C for 1 min (ramp rate of 2 °C/s) with a final inactivation step of 98 °C for 5 min. Following amplification, droplets were analyzed with a QX200 droplet reader and QuantaSoft vl.6.6 software (Bio-Rad). Relative transcript levels were expressed as a ratio of *Kcnb1* to *Tbp* with WT normalized to 1 and included 14–16 biological replicates per genotype. Normality of transcript and protein expression was assessed by D’Agostino & Pearson test and statistical comparison between groups was made using the nonparametric test Kruskal-Wallis with Dunn’s post-hoc comparisons (GraphPad Prism v9.4.1, Graph Pad Software, San Diego, CA). Both assays lacked detectable signal in genomic, no-RT and no template controls.

### 2.4 Immunoblotting

Forebrain P3 membrane protein fractions were isolated from male and female WT, *Kcnb1^C/+^* and *Kcnb1*^C/C^ mice at P58-83. Membrane fractions (50 μg) were separated on a 7.5% SDS-PAGE gel and transferred to nitrocellulose. Blots were probed with anti-KV2.1 (K89/34) and anti-mortalin/GRP75 antibodies (Table 2). Alexa Fluor 790 goat anti-mouse antibodies (Table 2) were used to detect signal on an Odyssey imaging system (LI-COR). Relative protein levels were determined by densitometry using Image Studio (LI-COR) and expressed as a ratio of KV2.1 to GRP75 with WT normalized to 1 and included 7-14 biological replicates per genotype.

**Table 2.**
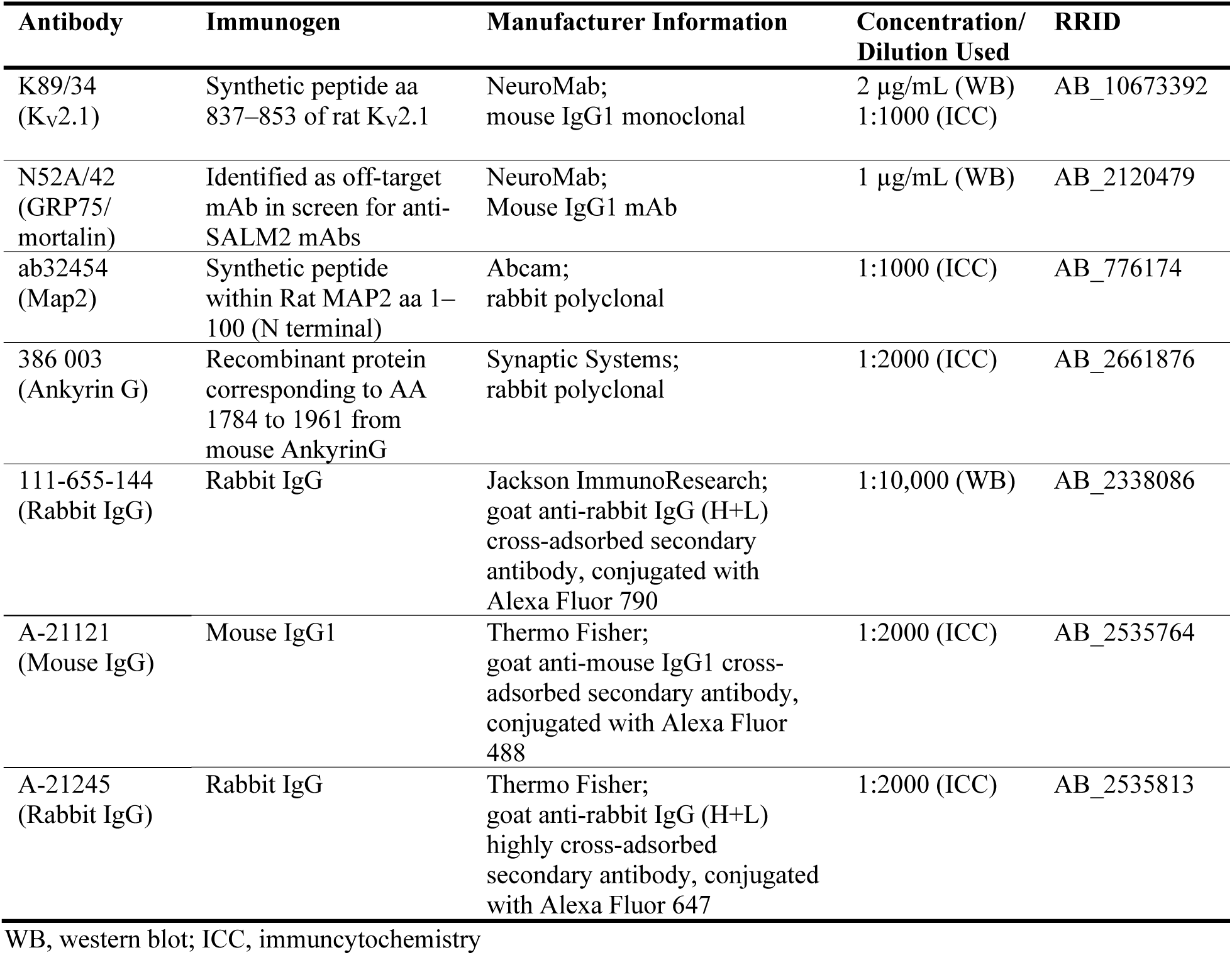
List of Antibodies

Normality of transcript and protein expression was assessed by D’Agostino & Pearson test and statistical comparison between groups was made using the nonparametric test Kruskal-Wallis with Dunn’s post-hoc comparisons (GraphPad Prism).

### 2.5 Primary neuron cultures

P0-1 pups were rapidly genotyped using the RFLP genotyping assay as described above. Hippocampal neurons were harvested and plated on poly-D-lysine-coated coverslips (GG-12-15-PDL; Neuvitro, Vancouver WA) at a density of 0.25-0.30x10e^6^ cells per well, maintained in Neurobasal medium supplemented with B-27 and Culture One (17504044 and A3320201; Gibco, Waltham, MA), with weekly half-volume media changes for 2-3 weeks. At least 3 independent cultures of 2 to 3 mice of each genotype were used for immunocytochemistry.

### 2.6 Immunocytochemistry and image analyses

At DIV18-21, coverslips were fixed using Cytofix/Cytoperm (554714; BD biosciences, San Jose CA) containing 4% sucrose (w/v) followed by additional permeabilization with 0.25% Triton X100 when necessary For Ankyrin G (AnkG) staining, two additional washes with 0.5% CHAPS (C3023; Sigma) in PermWash/PBS-T (PBS+tween-20) for 5 min were conducted. Coverslips were blocked with 10% normal goat serum for 30 mins. Incubations with primary antibodies for KV2.1 and Map2 or AnkG (Table 2) was performed in PermWash/PBS-T overnight at room temperature. Coverslips were then incubated with secondary antibodies (Table 2) diluted in PermWash/PBS-T + 10% NGS for 1-2h, followed by DAPI staining for 10 minutes. Coverslips were then mounted on glass slides using prolong gold antifade reagent (P36934; Invitrogen).

Images were acquired using a Nikon W1 spinning disk confocal and Hamamatsu camera in the Center for Advanced Imaging at Northwestern University. ND2 files were processed and analyzed using NIS elements software (Nikon). Images were identically processed in Adobe Photoshop (v23.2.1) for figure production.

Approximately 15 to 23 axon initial segment (AIS) measurements were taken from 10 to 12 randomly selected images at 60x magnification. For each genotype, 3 to 4 coverslips were evaluated.

### 2.7 Open field assay

Baseline locomotor activity was measured in P70-91 male and female WT, *Kcnb1^C/+^* and *Kcnb1*^C/C^ using an open field assay. Male and female mice were tested separately with at least a 1 h delay between sessions. Prior to behavioral testing, mice were acclimated in the behavior suite with white noise for 1 h. Each mouse was placed at the center of the open field arena (46cm x 46 cm) and video monitored for 30 min. Video records were analyzed offline using Ethovision XT software (Noldus Information Technology, Leesburg, VA, USA) by a reviewer blinded to genotype. Distance traveled, number of zone transitions and % time spent in center of arena for each trial were compared with one or two-way ANOVA with Tukey’s post hoc comparisons. No difference in sex was detected, therefore groups were by collapsed across sex (n=14-17 mice per genotype).

### 2.8 Seizure induction

#### 2.8.1 Flurothyl seizure induction

Susceptibility to seizures induced by the chemoconvulsant flurothyl (Bis(2,2,2-trifluoroethyl) ether, Sigma-Aldrich, St. Louis, MO) was tested in male and female WT, *Kcnb1^C/+^* and *Kcnb1*^C/C^ at P72-90 as previously described (Echevarria-Cooper et al., 2022; Hawkins et al., 2021).

Briefly, mice were placed in a Plexiglas chamber (2.2 L) and flurothyl was introduced using a syringe pump (20 uL/min) and allowed to volatilize. Latencies to first myoclonic jerk, generalized tonic-clonic seizure (GTCS) with loss of posture, and time interval between these phases were compared using Kruskal-Wallis with Dunn’s post-hoc comparisons. No difference in sex was detected, therefore groups were collapsed across sex (n=33-40 mice per genotype).

#### 2.8.2 Kainic Acid seizure induction

Susceptibility to seizures induced by the chemoconvulsant KA (kainic acid, Cat #0222, Tocris Bioscience, Minneapolis, MN) was tested in WT, *Kcnb1^C/+^* and *Kcnb1*^C/C^ mice at P41-50. KA dissolved in saline was administered by intraperitoneal injection (25 mg/kg) and mice were video recorded for 2 hours. Videos were scored offline by reviewers blinded to genotype using a modified Racine scale (Racine, 1972) (1-behavioral arrest; 2-forelimb and/or Straub tail, facial automatisms; 3-automatisms, including repetitive scratching, circling, forelimb clonus without falling; 4-forelimb clonus with rearing and/or falling, barrel roll; 5-repetition of stage 4; 6-generalized tonic-clonic seizure, wild running and/or jumping; 7-death). Latencies to the first occurrence of each stage and the highest seizure stage reached within 5 minutes bins were determined from video records by reviewers blinded to genotype. Latency to death following KA injection was compared by log-rank Mantel-Cox time to event analysis. Severity within time bins was compared between mutant alleles and WT by two-way ANOVA with Tukey’s post hoc comparisons. No difference in sex was identified, therefore groups were collapsed across sex (n=12-19 mice per genotype).

### 2.9 Video-EEG Monitoring

At 19-21 weeks of age, male and female WT, *Kcnb1^C/+^* and *Kcnb1*^C/C^ mice were implanted with EEG headmounts (8201, Pinnacle Technology, Lawrence, KS) under ketamine/xylazine anesthesia. Headmounts with four stainless steel screws that served as cortical surface electrodes were affixed to the skull with glass ionomer cement. Anterior screw electrodes were 0.5–1 mm anterior to bregma and 1 mm lateral from the midline. Posterior screws were 4.5–5 mm posterior to bregma and 1 mm lateral from the midline. EEG1 represents recordings from right posterior to left posterior (interelectrode distance ≈2 mm) and EEG2 represents recordings from right anterior to left posterior (interelectrode distance ≈5 mm). The left anterior screw served as the ground connection. Following at least 48 h of recovery, tethered EEG and video data were continuously collected for 7 to 14 days from freely moving mice with Sirenia acquisition software (Pinnacle Technology). Individual files were packeted into approximately 24-to-96-hour segments. EEG records were assigned a randomly generated code to blind reviewers to genotype during analysis. One file per mouse was chosen for manual review, and each file reviewed had a total average EEG baseline <75 µV with a low artifact incidence. On average, 67.5 hours of EEG data were analyzed from each subject (WT: 45-91 h/mouse (n = 4 mice); *Kcnb1^C/+^*: 47-73 h/mouse (n = 6 mice); *Kcnb1^C/C^*: 70-74 h/mouse (n = 5 mice)). EEG data between 0.5 and 200 Hz were acquired at a sampling rate of 400 Hz. Raw data was notch filtered around 60 and 120 Hz prior to analysis. Video-EEG records were manually reviewed for electrographic seizures (≥2x baseline; ≥10 sec; evolution in frequency and amplitude) and epileptiform discharges, including spike trains. Spike trains were defined as having regular, rhythmic sharp spike and slow wave components with ≥1 SWD per second continuing for ≥5 seconds, between 1-2.5 Hz frequencies.

### 2.10 EEG spectral analysis

To quantify baseline EEG activity and generate spectral arrays from Figures 5 and 6, we used a Fast Fourier Transform (FFT) algorithm with an FFT size of 1024, 93.75% overlap window and Hann-Cosine-Bell fit, which generated approximately 330 frequency data points between 0-128 Hz. After visual inspection to exclude artifacts, we used Lab Chart v.8.1.19 (DSI) for spectral analysis and max power frequency data. Power analysis was performed on EEG baseline recordings for each file selected. PSD values were generated from 13-h segments during the light cycle (1-3 PSDs/mouse, 9-11 PSDs/genotype) and a 9-h segment during the dark cycle (2-4 PSDs/mouse, 13-17 PSDs/genotype). All PSD frequencies (per genotype) were analyzed for outliers by ROUTs. Statistical analysis was performed for each frequency band, delta (0.1–4 Hz), theta (5–8 Hz), alpha (8–13 Hz) and beta (14–29 Hz), using Two-way ANOVA with Tukey’s multiple comparison test. Maximum power frequency was calculated for each light or dark cycle segment reviewed (Lab Chart v.8.1.19). Light and dark values for each genotype were compared using Student’s T-test or Mann-Whitney. There were no significant differences between light or dark; therefore, all light and dark maximum frequency values were combined for each genotype, with 22-24 values per genotype.

### 2.11 Statistical analysis

Table 3 summarizes statistical tests used for all comparisons along with computed values. Values for post-hoc tests are reported in the results and figure legends. No significant differences were detected between sexes on reported measures; thus, groups were collapsed across sex variables.

**Table 3.**
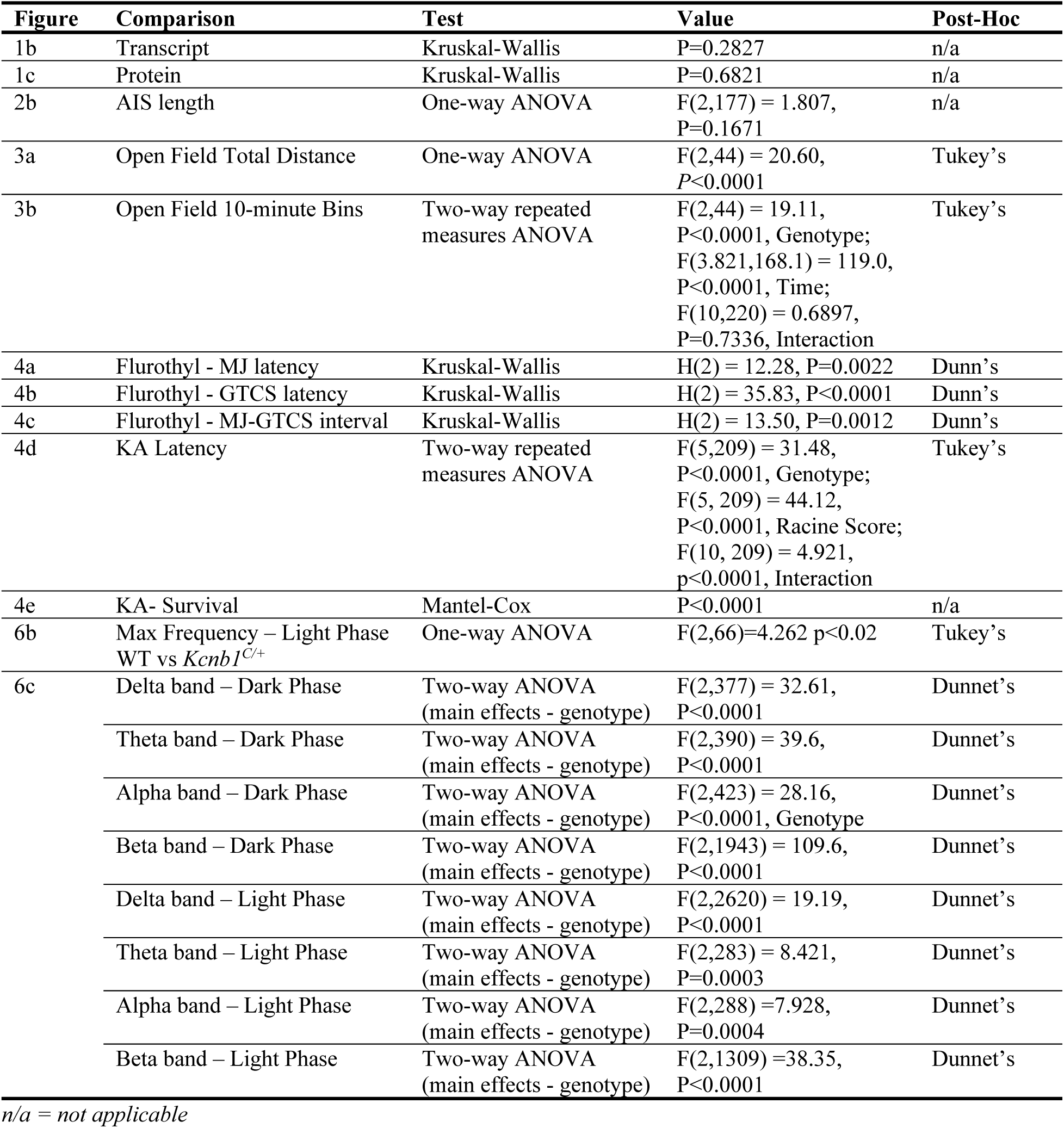
Summary of Statistical Comparisons

## 3. Results

Previous functional studies showed altered voltage-dependence of KV2.1 channels incorporating the R306C variant (Kang et al., 2019; Saitsu et al., 2015). In transfected CHO-K1 cells, despite normal cell surface expression of KV2.1, R306C effects ranged from complete loss of delayed rectifier potassium currents when singly expressed to partial loss of function and altered voltage-dependence when co-expressed with WT to approximate the heterozygous condition (Kang et al., 2019). Transfection into cultured cortical pyramidal neurons resulted in lower sensitivity and cooperativity of the voltage sensor and severely impaired repetitive firing (Saitsu et al., 2015).

These effects on channel function are unique compared to other variants modeled in mouse to date, including the dominant-negative pore variant *Kcnb1^G379R^*, the trafficking defective variant *Kcnb1^R312H^*, *Kcnb1* knockout mice (*Kcnb1^-/-^*), or premature termination codon *Kcnb1*^fs^ mice (Bortolami et al., 2022; Hawkins et al., 2021; Speca et al., 2014). Furthermore, the R306C variant has a high rate of recurrence in patient cohorts. Taken together, these observations made R306C a high priority variant for *in vivo* modeling.

### 3.1 Generation and initial characterization of Kcnb1^R306C^ mice

*Kcnb1^R306C^* mice on the C57BL/6J inbred strain were generated using CRISPR/Cas9 to introduce the modification of arginine 306 (same codon number in human and mouse) by HDR. Sequencing chromatograms of *Kcnb1* genomic PCR products showing the WT and R306C variant alleles are shown in Figure 1A. Expression analyses of bulk forebrain tissue revealed no significant differences in the *Kcnb1* transcript or KV2.1 protein expression across the three genotypes: WT, heterozygous *Kcnb1^R306C/+^* (*Kcnb1^C/+^*) and homozygous *Kcnb1^R306C/R306C^* (*Kcnb1*^C/C^) (Fig. 1B-D; *p* > 0.2, Kruskal-Wallis). Thus, the R306C variant does not affect KV2.1 expression levels, consistent with prior studies in CHO-K1 cells (Kang et al., 2019).

**Figure 1.**
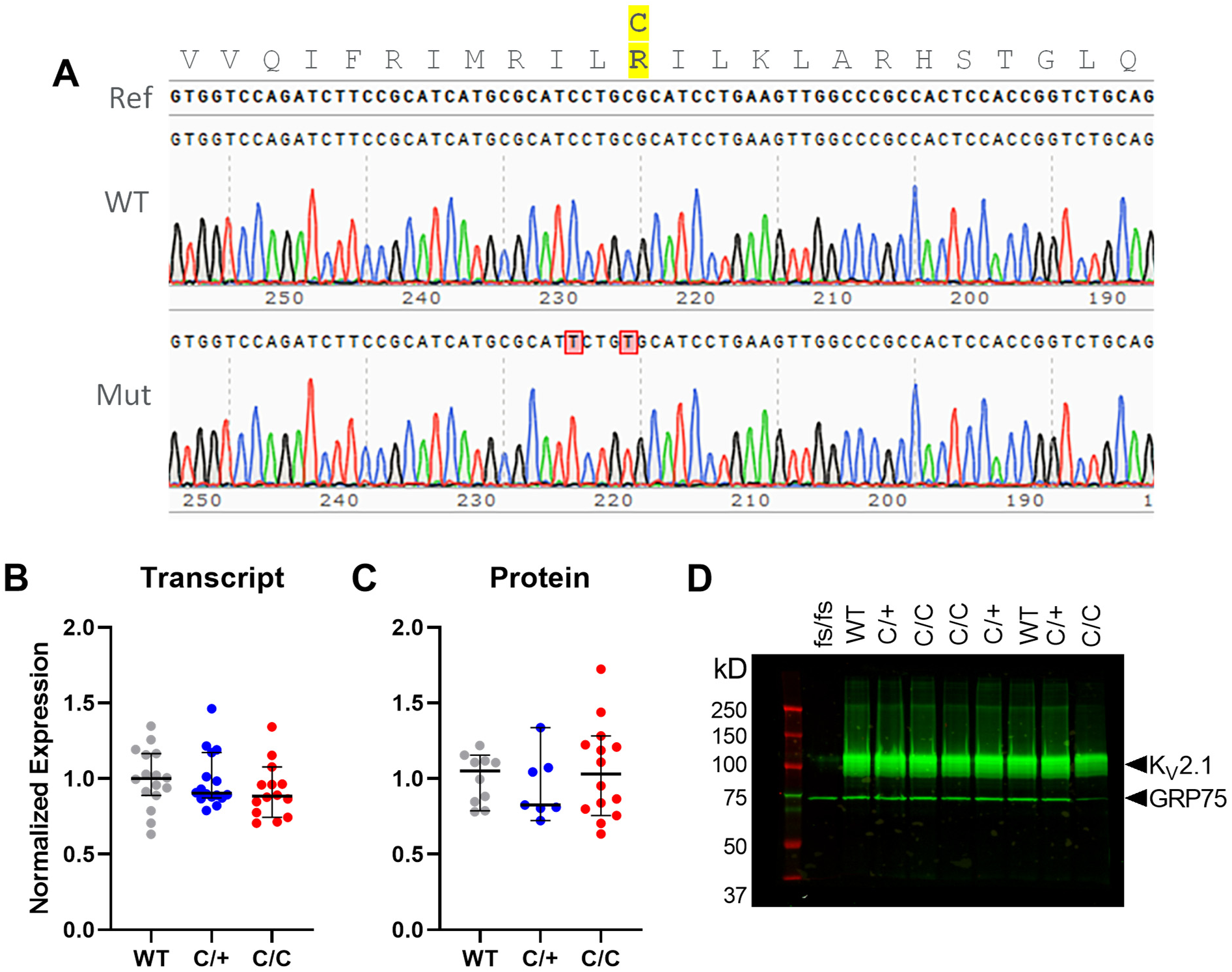
Characterization of *Kcnb1*-p.R306C variant allele. **(A)** Chromatogram confirmation of *Kcnb1* R306C genomic PCR product. Genomic PCR products were TOPO cloned and Sanger sequenced. The top trace represents a WT allele with 100% identity to the mouse C57BL/6J reference genome (GRCm39). The bottom trace shows a mutated allele (Mut) that encodes p.R306C, as well as a silent mutation to destroy the PAM site (p.I304=). **(B-C)** Transcript (B) and protein (C) expression levels did not differ between genotypes genotypes (p=0.28 and 0.68 respectively, Kruskal-Wallis). Symbols represent samples from individual mice, line represents median and error bars represent 95% confidence interval. **(D)** A representative western blot of forebrain membrane proteins probed for Kv2.1 protein and GRP75 as a normalization control show no difference. *Kcnb1*^fs/fs^ that results in truncated protein with absent epitope was used as a negative control.

### 3.2 Normal expression and localization of KV2.1 in R306C cultured hippocampal neurons

To assess the impact of the *Kcnb1^R306C^* variant on KV2.1 expression in neurons, we performed immunolabeling of cultured hippocampal neurons (DIV16-18) isolated from WT, *Kcnb1^C/+^*and *Kcnb1*^C/C^ mice.

The *Kcnb1^R306C^* variant had no overt effect on expression or subcellular localization of KV2.1. Robust clustering in the soma and proximal processes was present across all genotypes, indicating that the R306C variant does not impair localization or cluster formation (Fig. 2A). Co-immunolabeling with MAP2 or Ankyrin-G was consistent with KV2.1 localization in proximal dendrites and AIS, respectively, across all genotypes (Fig. 2A). One hallmark of KV2.1 expression in neurons is its early and robust expression in the proximal AIS, a critical site for regulation of neuronal polarity. Therefore, we measured AIS length to determine if *Kcnb1*-p.R306C affected AIS maturation. AIS length measurements were not different across genotypes (Fig. 2B) (F(2,177) = 1.807, *p* = 0.1671, one-way ANOVA).

**Figure 2.**
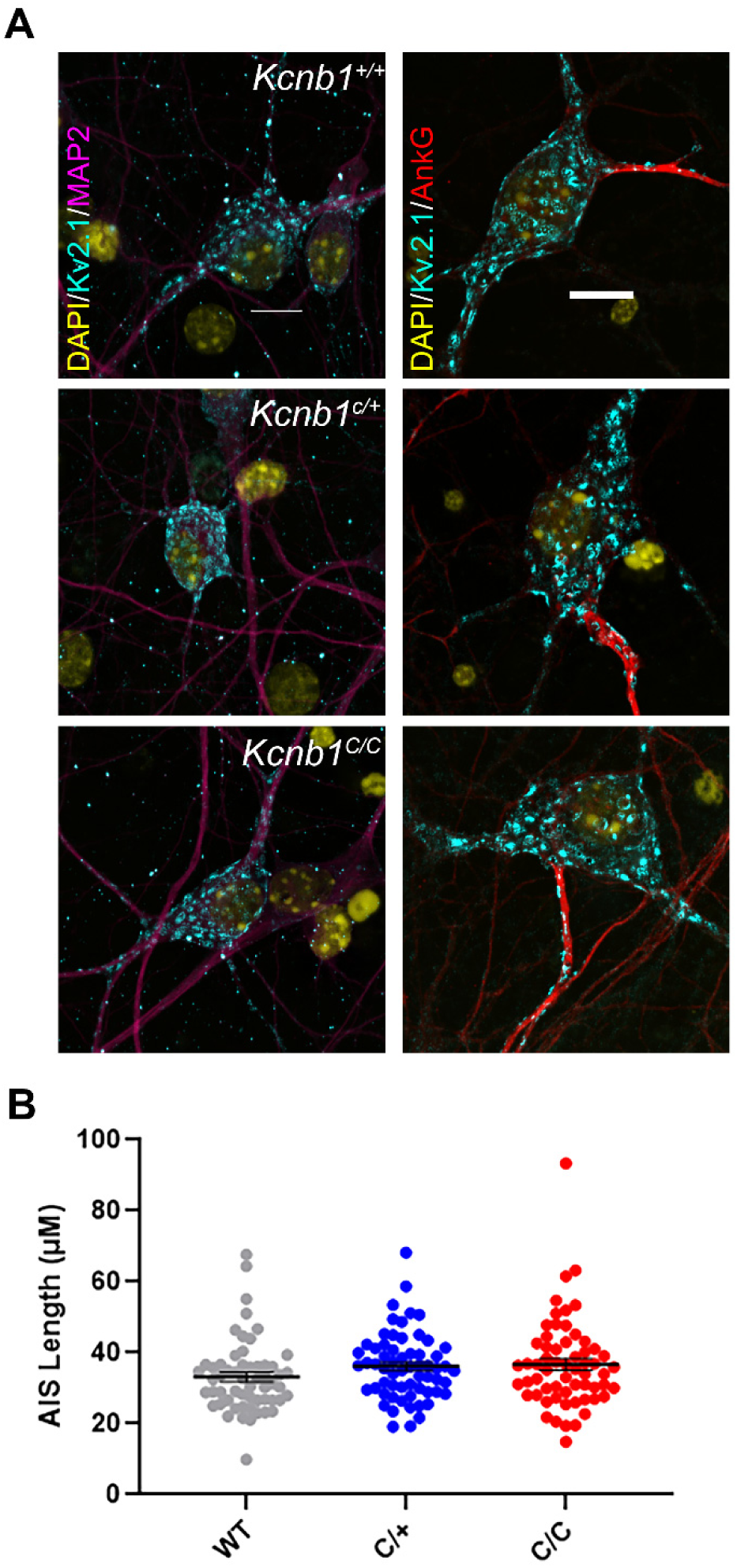
Subcellular localization Kv2.1 in cultured hippocampal neurons. **(A)** Cultured hippocampal neurons were immuno-labeled with Kv2.1 and neuronal processes (MAP2 for dendrites and AnkG for axon initial segment); 40x magnification for MAP2 and 60x for AnkG staining. (**B)** Length measurements of AIS were not different across different genotypes (*p* = 0.1671, one-way ANOVA; n = 57-62 per genotype).

### 3.3 Locomotor hyperactivity in Kcnb1^R306C^ mice

Differences in locomotor activity levels between *Kcnb1^R306C^* and WT mice was evident in their home cages during routine husbandry and was reminiscent of the profound hyperactivity observed in *Kcnb1^G379R^* and *Kcnb1^-/-^* mice (Hawkins et al., 2021; Speca et al., 2014). To quantify this effect, we measured baseline locomotor activity in a 30-minute open field assay. Both *Kcnb1^C/+^* and *Kcnb1^C/C^* mice traveled farther in the open field than WT littermate controls (F(2,44) = 20.6, *p* < 0.0001, one-way ANOVA). WT mice traveled an average distance of 121.4 ± 4.1 m, while *Kcnb1^C/+^* and *Kcnb1^C/C^* traveled 155.1 ± 5.4 and 164.7 ± 5.4 m, respectively (Fig. 3A,E). Distance traveled was consistently elevated over the course of the 30-minute period in *Kcnb1^C/+^*and *Kcnb1^C/C^* mice compared to WT, while all genotypes showed similar short-term habituation (Fig. 3B). The number of zone transitions and time spent in the arena center were assessed as a rudimentary evaluation for anxiety within the open field setting. The total number of zone transitions was affected by *Kcnb1* genotype (F(2,44) = 4.86 *p* < 0.02; one-way ANOVA). *Kcnb1^C/+^* mice crossed the testing arena an average of 207.4 ± 13.5 times in 30 minutes (*p* < 0.02), while WT and *Kcnb1^C/C^* mice crossed an average of 150.6 ± 10.8 times and 192.5 ± 16.6, respectively (Fig. 3C,E). However, there was no difference between genotypes for the percentage of time spent in the center of the arena (Fig. 3D).

**Figure 3.**
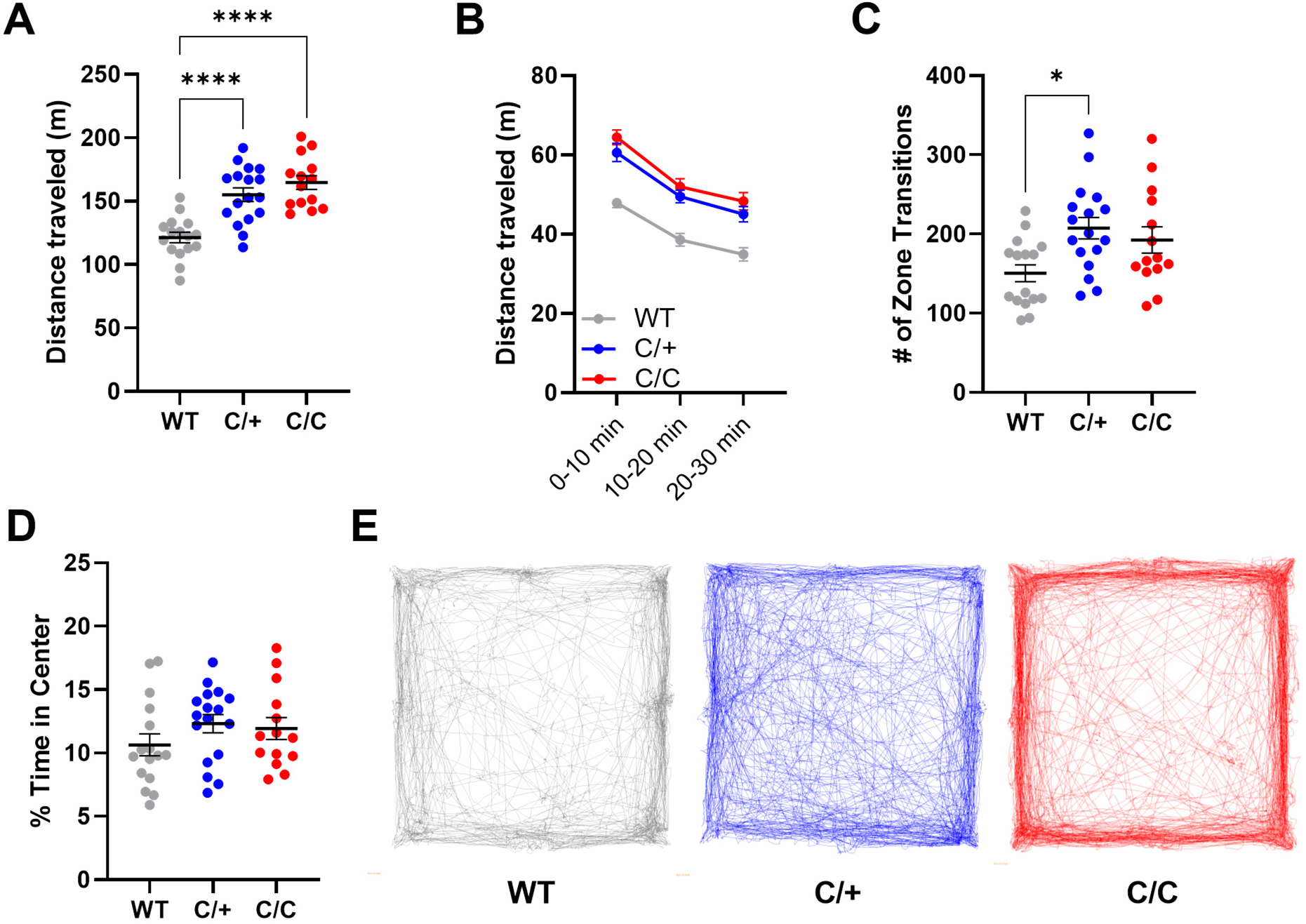
*Kcnb1^R306C^* mice have elevated exploratory locomotion in open field assay. **(A)** Total distance traveled in 30 minutes was affected by genotype (F(2,44) = 20.6, *p* < 0.0001; one-way ANOVA; \*\*\*\**p* < 0.0001, Tukey’s). (**B)** Locomotion assessed in 10-minute bins across the 30-minute session. *Kcnb1^C/+^* and *Kcnb1^C/C^*mice had elevated distance at all time points compared to WT (F(2,44) = 19.11, *p* < 0.0001; Two-way repeated measures ANOVA, Genotype). All groups showed similar habituation, with locomotion slowing over time and no significant genotype-by-time interaction (see Table 3). **(C)** Total number of zone transitions was affected by genotype (F(2,44) = 4.86, *p* < 0.02; one-way ANOVA; \**p* < 0.02, Tukey’s). (**D**) Time spent in the center of area was not affected by genotype (F(2,44) = 1.201, *p* > 0.31). (**E**) Representative examples of open field paths for WT, *Kcnb1^C/+^*and *Kcnb1^C/C^* mice. For A, C and D, symbols represent individual mice and horizontal lines represent mean. Symbols in B represent group means. Error bars represent SEM (A-D).

### 3.4 Altered seizure susceptibility in Kcnb1^R306C^ mice

Previous studies showed enhanced seizure susceptibility in *Kcnb1^G379R^* and *Kcnb1^-/-^*knockout mice (Hawkins et al., 2021). In order to compare with other *Kcnb1* mouse models, we asked how the R306C variant affects seizure susceptibility using two chemoconvulsants, flurothyl and KA. First, we used the GABAA antagonist flurothyl to induce a stereotyped progression that begins with a myoclonic jerk (MJ) as the first seizure sign and progresses to a GTCS. Latency for flurothyl-induced seizures was affected differently by mutant allele dosage. *Kcnb1^C/+^*mice exhibited longer latencies to both the MJ (127 seconds 95% CI [115, 141]) and GTCS, (188 seconds 95% CI [170, 205]), compared to WT mice (MJ: 113 seconds 95% CI [101, 121]; GTCS: 173 seconds 95% CI [158, 179]) (Fig. 4A-B). In contrast, *Kcnb1^C/C^* mice showed no difference in MJ latency relative to WT, but had a ≍15% shorter latency shorter latency to the GTCS (132 seconds 95% CI [128 146]), (Fig. 4A-B). Comparison of the time interval between MJ and GTCS showed that *Kcnb1^C/C^* mice progressed quickly between the stages, with a median time of 27 seconds 95% CI [17-50], while WT and *Kcnb1^C/+^* exhibited similar median latencies of 58 seconds 95% CI [45-68] and 53 seconds 95% CI [40-64], respectively (Fig. 4C).

**Figure 4.**
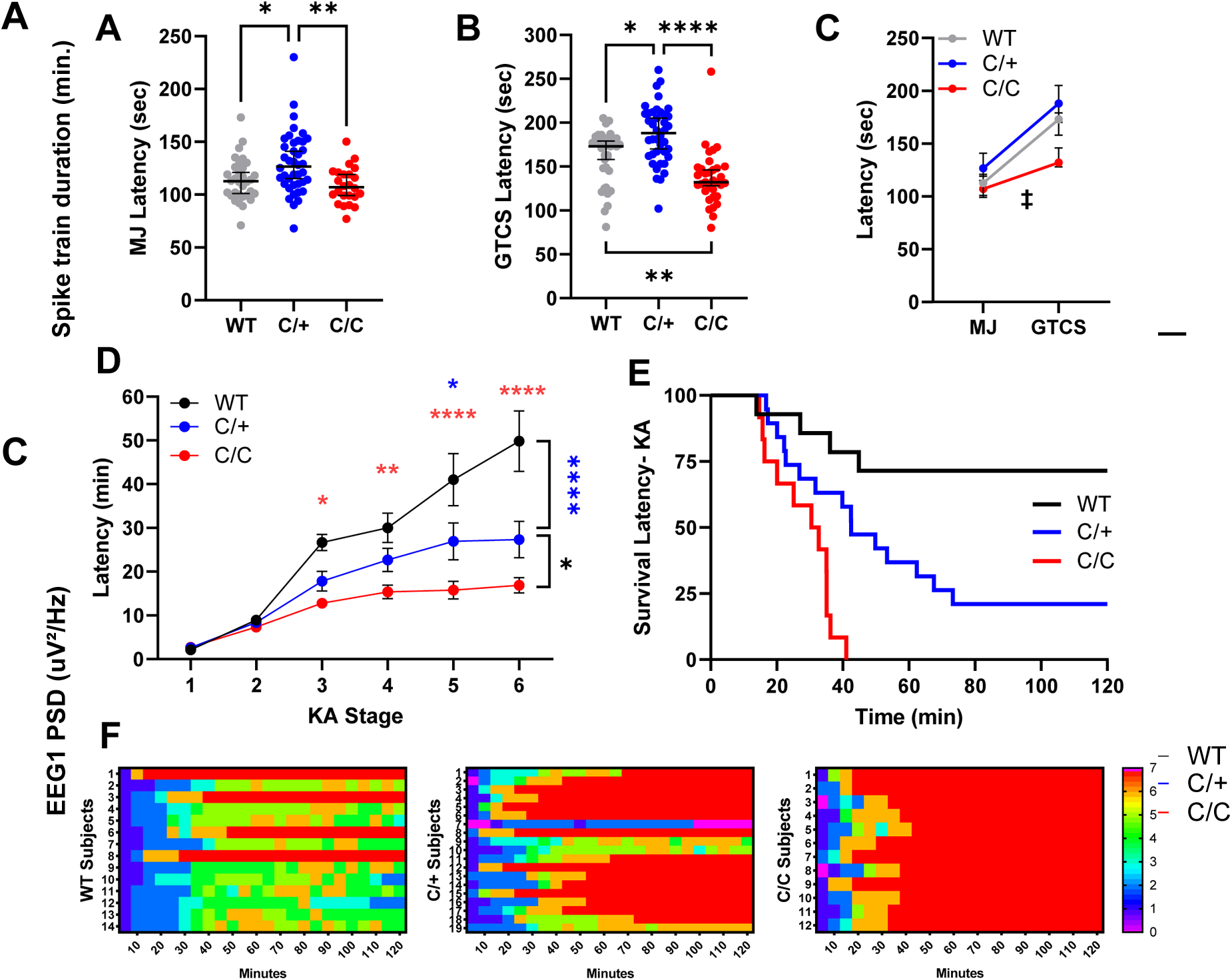
Altered susceptibility to seizures induced by chemoconvulsants in *Kcnb1^C/+^*and *Kcnb1^C/C^* mice. **(A)** Latency to the first flurothyl-induced MJ was affected by *Kcnb1* genotype (Kruskal-Wallis H(2) = 12.28, *p* < 0.003; \**p* < 0.04 and \*\**p* < 0.004, Dunn’s). **(B)** Latency to flurothyl-induced GTCS was affected by *Kcnb1* genotype (Kruskal-Wallis H(2) = 35.83, *p* < 0.0001). *Kcnb1^C/+^* mice exhibited the longest median latency to GTCS, while *Kcnb1^C/C^* mice had the shortest latency to GTCS (\**p* < 0.03, \*\**p* < 0.003, \*\*\*\**p* < 0.0001, Dunn’s). For A-B, symbols represent individual mice, horizontal line represents median and error bars are 95% CI. **(C)** Progression from MJ to GTCS differed between genotypes (Kruskal-Wallis H(2) = 13.50, *p* < 0.002). *Kcnb1^C/C^* mice progressed the fastest between stages compared to WT (‡ *p* < 0.005) and *Kcnb1^C/+^* (‡ *p* < 0.003). **(D)** Progression of seizure stages scored on a modified Racine scale post KA administration. *Kcnb1^C/C^* mice progressed faster through stages 3-6 compared to WT and to stage 6 compared to *Kcnb1^C/+^* (WT v. *Kcnb1^C/C^*Red \**p* < 0.02 \*\**p* < 0.008, \*\*\*\**p* < 0.0001; *Kcnb1^C/+^* v. *Kcnb1^C/C^* Black \**p* < 0.03). *Kcnb1^C/+^* mice progressed faster to stages 5 and 6 compared to WT (WT v. *Kcnb1^C/+^* Blue \**p* < 0.02, \*\*\*\**p* < 0.0001). *P* values determined by Tukey’s with Two-way ANOVA (see Table 3). Symbols represent the average latency of n = 12-19 mice per genotype. Error bars represent SEM. **(E)** Survival after KA administration was affected by *Kcnb1* genotype. 100% of *Kcnb1^C/C^* and 75% of *Kcnb1^C/+^* mice did not survive following KA administration, compared to 25% of WT mice (LogRank Mantel-Cox *p* < 0.0001). **(F)** Heat map depicting individual level seizure severity post KA administration organized by genotype. Y axis represents individual mice and X axis represents 5 min bins.

Next, we evaluated susceptibility to seizures induced by the glutamatergic agonist KA in a separate cohort of mice. Seizure intensity following KA administration was assessed over a 2-hour period using a modified Racine scale, scoring for latency to first occurrence of each stage and for the highest stage reached within 5-minute bins (Fig. 4D-F). There were no differences in latency to stages 1 or 2 between any genotype. However, for stages 3-7, *Kcnb1^C/C^*mice had a shorter average latency compared to WT and was quicker to reach stage 6 compared to *Kcnb1^C/+^*. (Fig. 4D). *Kcnb1^C/+^* mice had shorter latencies to stages 5 and 6 faster compared to WT mice (Fig. 4D). Comparison of the survival rates of WT, *Kcnb1^C/+^*and *Kcnb1^C/C^* after KA administration revealed that all *Kcnb1^C/C^* and ≍75% of *Kcnb1^C/+^* died within 2 hours following KA administration, while only ≍25% of WT mice died (Fig. 4E-F). The median latency to death was 31.6 min in *Kcnb1^C/C^* and 42.5 min for *Kcnb1^C/+^* mice, while ≍75% of WT mice survived for at least 2-h post-KA (Fig. 4E-F).

### 3.5 EEG Abnormalities in Kcnb1^R306C^ mice

Adult *Kcnb1^C/C^* mice were sporadically observed exhibiting spontaneous GTCS in their home cages (Supplemental Video S1). To systematically evaluate electrographic events and quantify EEG properties, we collected synchronized video-EEG data from WT, *Kcnb1^C/+^*and *Kcnb1^C/C^* mice at ≈20 weeks of age. Video-EEG was continually recorded for 7-14 days. Manual review of EEGs revealed a single spontaneous generalized seizure in a *Kcnb1^C/+^*mouse (Fig. 5A; Supplemental Video S2), as well as a single generalized seizure in a *Kcnb1^C/C^* mouse. Low GTCS incidence was not unexpected, as previous work from our lab showed that other *Kcnb1* mutant lines had relatively low GTCS frequency (Hawkins et al., 2021; Speca et al., 2014). Furthermore, GTCS witnessed during routine handling or husbandry were infrequent.

**Figure 5.**
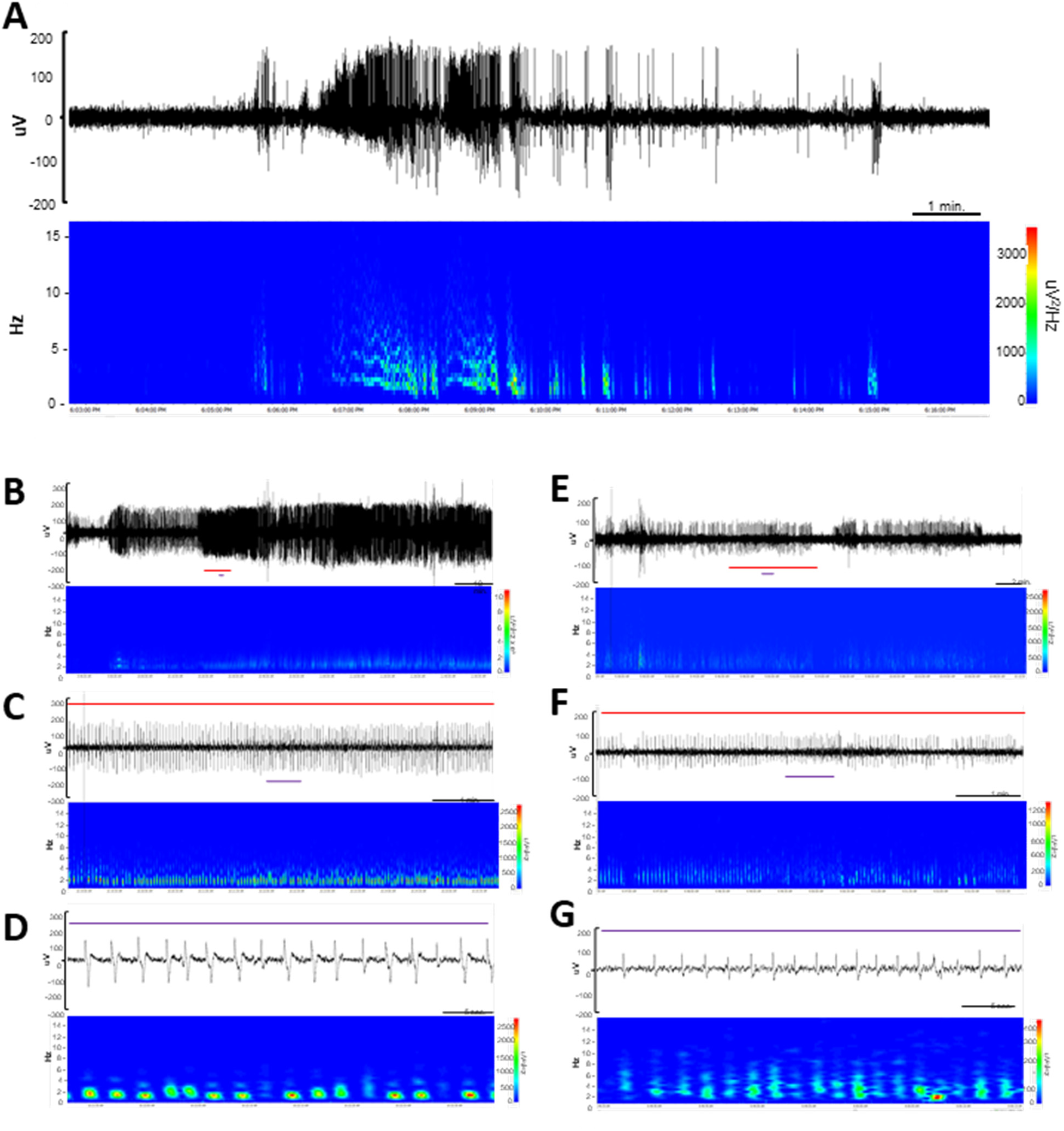
EEG abnormalities in *Kcnb1^C/+^* and *Kcnb1^C/C^* mice. **(A)** EEG trace of the single GTC seizure identified in a *Kcnb1^C/+^* mouse with corresponding spectral density array. Corresponding with the electrographic seizure, the *Kcnb1^C/+^* mouse displayed head bobbing and facial automatisms. All observable behaviors concluded when the baseline EEG activity returned to normal. One additional GTCS was detected in a *Kcnb1^C/C^*mouse with similar electrographic and behavioral manifestations. **(B-D)** Top trace represents a two-hour segment of continuous spike-wave discharges identified in a *Kcnb1^C/+^* mouse with corresponding spectral density array (B). Red line represents a ∼7-minute segment expanded in panel C and the purple line represents a ∼40 second segment expanded in panel D. (**E-G)** Top trace represents a ∼30-minute segment of continuous spike-wave discharges identified in a *Kcnb1^C/C^*mouse with corresponding spectral density array (E). Red line represents a ∼7-minute trace expanded in panel F and purple line represents a ∼ 40 second expanded in panel G.

Beyond the rare spontaneous GTCS events, EEG was markedly abnormal in *Kcnb1^R306C^* mice. Both *Kcnb1^C/+^*and *Kcnb1^C/C^* mice displayed recurrent trains of slow spike-wave complexes of varying durations on EEG (Fig. 5B-G, 6A). To quantify these observations, we performed focused analysis of approximately 67.5 hours of EEG records with low artifact and total average EEG baseline < 75 uV (WT: range of 45-91 hrs/mouse (n = 4); *Kcnb1^C/+^*: range of 47-73 hrs/mouse (n = 6); *Kcnb1^C/C^*: range of 70-74 hrs/mouse (n = 5)). Within these records, we first assessed frequency and duration of spike trains by manual review of EEG and video records.

Spike-wave trains were more often identified during periods of rest, but many did occur during wakefulness and coincided with circling or behavioral arrest with head bobbing (Fig. 5B-G). In *Kcnb1^C/+^* recordings, 4 of 6 mice collectively experienced 69 instances of spike wave trains ranging from 6 seconds to 140 minutes (Fig. 5B-D, 6A). Spectral analysis during these runs demonstrated elevated spectral density units (µV^2^/Hz) in the frequency range of 1-4 Hz (Fig. 5B-D). In *Kcnb1^C/C^* recordings, 3 of 5 mice collectively experienced 45 instances, ranging from 19 econds to 108 minutes, again with elevated spectral density units (µV^2^/Hz) in the frequency range of 1-4 Hz (Fig. 5E-G, 6A). Similar spike trains were never observed in recordings from WT mice.

**Figure 6.**
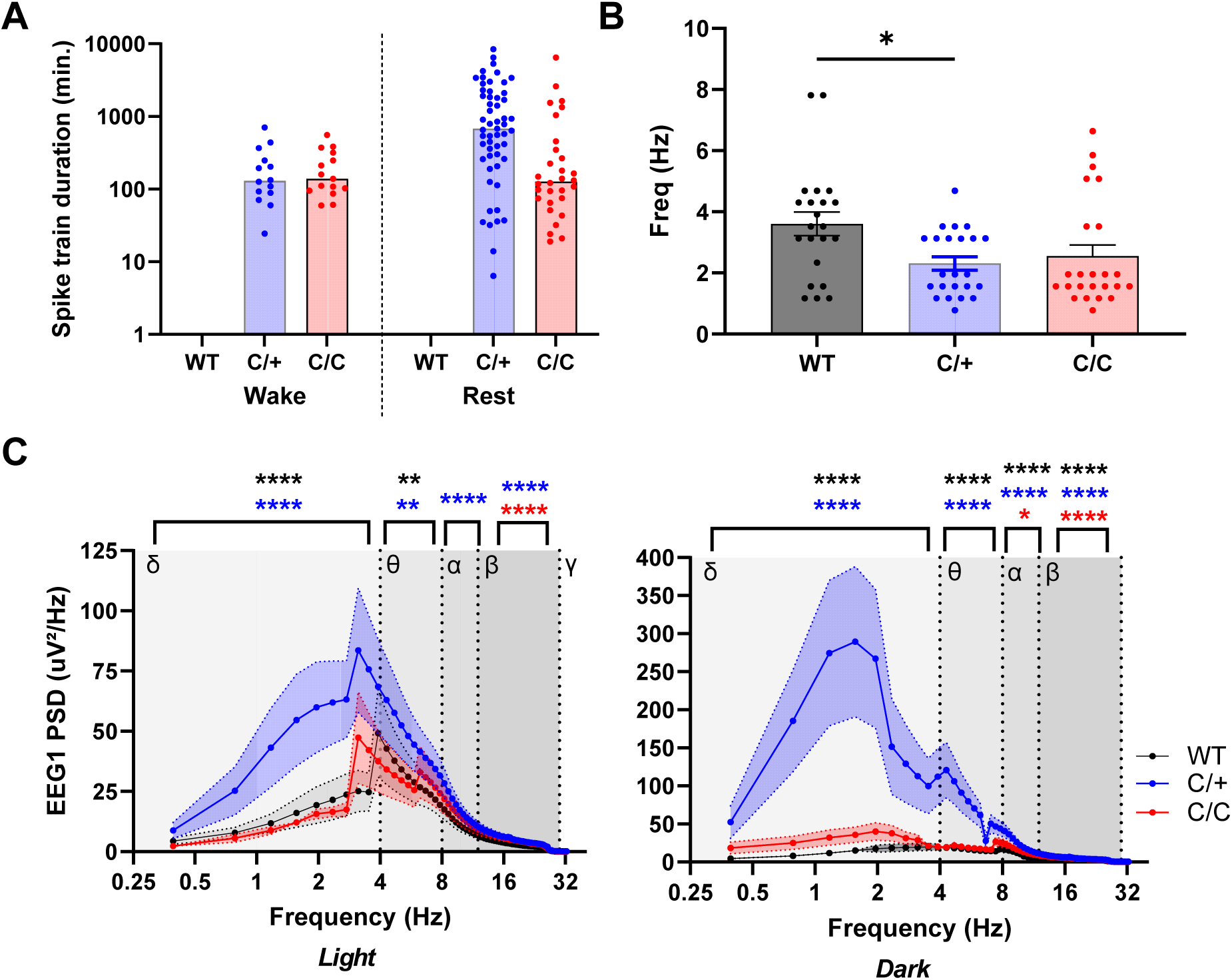
Abnormal EEG power in *Kcnb1^C/+^* and *Kcnb1^C/C^* mice. **(A)** Spike train frequency and durations for *Kcnb1^R306C^* mice. During manual review of EEG records (45-91 hrs per mouse), four *Kcnb1^C/+^*(66%) and 3 *Kcnb1^C/C^* (60%) mice exhibited recurrent spike trains of various durations, while never detected in the EEG files reviewed from WT mice. Bars represent median length of spike trains. Symbols represent durations of individual trains. Left side shows trains during wakeful behavior and right side shows trains during restful behavior. **(B)** Max power frequency for WT, *Kcnb1^C/+^*and *Kcnb1^C/C^* mice. WT mice had a calculated average max power frequency of 3.6 ± 0.4 Hz, while *Kcnb1^C/+^*averaged 2.3 ± 0.2 Hz (\**p* < 0.03) and *Kcnb1^C/C^* averaged 2.6 ± 0.4. Bars represent mean and error bars SEM. Symbols represent individual values. No difference in max frequency was detected between light and dark cycles, therefore they were combined, resulting in 22-24 values per genotype. *P* values calculated by one-way ANOVA with Tukey’s. **(C)** Power spectra in *Kcnb1* mice. Power band comparisons for delta (0.4 - 4 Hz) theta (4 - 8 Hz), alpha (8 - 13 Hz) and beta (13 - 30 Hz) during light and dark phases. In the light phase, compared to WT, *Kcnb1^C/+^* mice had elevated PSD in the delta (*p* < 0.0001), theta (*p* < 0.002), alpha (*p* < 0.0003) and beta (*p* < 0.0001) frequency bands, while *Kcnb1^C/C^* mice had elevated beta PSD (*p* < 0.0001) compared to WT. In the dark phase, compared to WT, *Kcnb1^C/+^*had elevated PSD across all frequency bands (*p* < 0.0001), while *Kcnb1^C/C^*had elevated PSD in the alpha (*p* < 0.02) and beta (*p* < 0.0001) frequency bands. PSD represents average power from n = 4-6 mice per genotype, with 9-17 values per frequency. Symbols represent average frequency and error bars are SEM. Black asterisks represent significance between *Kcnb1^C/+^* and *Kcnb1^C/C^*, red asterisks represent significance between WT and *Kcnb1^C/C^* and blue asterisks represent significance between WT and *Kcnb1^C/+^*. *P* values calculated by two-way ANOVA with Tukey’s.

We also used quantitative EEG spectral analysis of as a proxy measurement of global (ictal and interictal) activity across the light (13-h segments) and dark cycles (9-h segments). Max power frequency did not differ between light and dark phases within genotype and was therefore collapsed across phase. Both *Kcnb1^C/+^* and *Kcnb1^C/C^* mice displayed high powered activity in the ∼2 Hz frequency range (2.3 ± 0.2, 2.6 ± 0.4, respectively), while WT had an average max power at ∼ 4 Hz (3.6 ± 0.4) (Figure 6B). This frequency difference is likely attributed to the recurrent, pathogenic SWDs found throughout *Kcnb1^C/+^*and *Kcnb1^C/C^* EEGs.

Examination of power spectral densities (PSD) between 0-30 Hz in the dark phase showed that compared to WT, *Kcnb1^C/+^*had elevated PSD across all frequency bands (*p* < 0.0001 for each band). In contrast, *Kcnb1^C/C^* had elevated PSD in the alpha (*p* < 0.02) and beta (*p* < 0.0001) frequency bands, while delta and theta PSDs were not significantly different (Fig. 6C; Table 3). Examination of PSD in the light phase revealed that *Kcnb1^C/+^* had elevated PSD in the delta (*p* < 0.0001), theta (*p* < 0.002), alpha (*p* < 0.0003) and beta (<0.0001) frequency bands, while *Kcnb1^C/C^* mice had elevated beta PSD (*p* < 0.0001) (Fig. 6C; Table 3). The observed alterations of PSD in *Kcnb1* mutant mice relative to WT are suggestive of an imbalance in excitation-inhibition homeostasis.

## 4. Discussion

*KCNB1*-p.R306C is the one of the most recurrent variants identified to date in individuals with *KCNB1* encephalopathy, and is associated with mild to severe global developmental delays, behavioral disorders, and a diverse spectrum of epilepsy that includes infantile spasms, GTC, myoclonic, tonic, focal, and absence seizures (Bar et al., 2020a; de Kovel et al., 2017; Kang et al., 2019; Marini et al., 2017; Saitsu et al., 2015). Previous *in vitro* characterization of R306C showed loss of voltage sensitivity and cooperativity of the voltage sensor and inhibition of repetitive firing, which can disrupt neuronal circuitry and result in clinical manifestations of developmental encephalopathy (Kang et al., 2019; Saitsu et al., 2015). We and others have previously generated and characterized mouse models of *Kcnb1* encephalopathy, including the missense variants *KCNB1*-p.G379R, *KCNB1*-p.R312H, as well as a *Kcnb1^-/-^* knock-out and frameshift alleles (Bortolami et al., 2022; Hawkins et al., 2021; Speca et al., 2014). The existing *Kcnb1* mouse models recapitulated *KCNB1* encephalopathy phenotypes, including spontaneous seizures, EEG abnormalities, learning deficits and hyperactivity (Bortolami et al., 2022; Hawkins et al., 2021; Speca et al., 2014). While *Kcnb1^G379R^*, *Kcnb1^R312H^*, and *Kcnb1* null and frameshift models are informative LoF alleles, the *Kcnb1*^R306C^ mutation represents a recurrent variant with different underlying mechanisms and biological impacts. Thus, this new model expands the allelic series of *Kcnb1* mouse models, which may help establish nuanced genotype-phenotype correlations.

Previous results from our laboratory demonstrated that the R306C variant exhibited normal protein expression level and cell-surface trafficking in CHO cells (Kang et al., 2019). Consistent with this, *Kcnb1^C/+^* and *Kcnb1^C/C^*mice did not have significant changes in transcript, or protein expression relative to WT (Fig. 1). Moreover, there was no difference in subcellular localization or clustering in *Kcnb1^C/+^* and *Kcnb1^C/C^* mice compared to WT (Fig. 2). In contrast, the *Kcnb1*^G379R^ and *Kcnb1^R312H^* mouse models had substantial, genotype-dosage dependent reduction in KV2.1 expression, as well as dominant-negative effects on KV2.1 macromolecular complexes (Bortolami et al., 2022; Hawkins et al., 2021). Thus, the *Kcnb1*^R306C^ phenotype is likely a direct result of defective voltage sensing rather than diminished protein expression and/or localization. This is an important distinction as voltage-sensing dysfunction may be amenable to different therapeutic approaches than diminished protein expression (Haq et al., 2022; Tian et al., 2022).

Both *Kcnb1*^C/+^ and *Kcnb1^C/C^*mice exhibited open field hyperactivity. This is consistent with previous reports of hyperactivity in *Kcnb1^G379R^* and *Kcnb1^-/-^* mice (Hawkins et al., 2021; Speca et al., 2014), as well as clinical reports of attention-deficit/hyperactivity disorder or hyperactivity with inattention in numerous cases of *KCNB1* encephalopathy (Bar et al., 2020a; de Kovel et al., 2017; Kang et al., 2019; Marini et al., 2017; Torkamani et al., 2014). There was no significant effect of R306C allele dosage on hyperactivity, in contrast to G379R that consistently had more severe neurobehavioral phenotypes in homozygotes compared to heterozygous littermates (Hawkins et al., 2021). This again supports the notion of differential effects of defective voltage sensing versus dominant negative in *Kcnb1* disease.

Seizure susceptibility was altered in *Kcnb1^R306C^*mice relative to WT, although there were differential effects depending on genotype and chemoconvulsant. In response to KA, we observed an allele dosage effect, with *Kcnb1^C/C^* mice being more severe than both *Kcnb1^C/+^* and WT, and *Kcnb1^C/+^* having intermediate sensitivity to KA. In contrast, latency to flurothyl-induced seizures was differentially modulated in *Kcnb1^C/+^* versus *Kcnb1^C/C^* mice. Latencies relative to WT were longer in *Kcnb1^C/+^* heterozygotes, while they were shorter in *Kcnb1^C/C^*homozygotes. The paradoxical relationship between genotype and seizure susceptibility in response to flurothyl suggests a fundamental difference between having mutant-only homotetramers in *Kcnb1^C/C^*mice versus *Kcnb1^C/+^* mice having a mixture of possible KV2.1 tetramer populations that may include WT-only and mutant-only homotetramers, as well as heterotetramers of WT and R306C with various stoichiometric ratios (Kang et al., 2019). When expressed as a homotetramer, R306C channels had complete loss-of-function. Consistent with this, the observed seizure susceptibility in *Kcnb1^C/C^*homozygous mice was similar to *Kcnb1^-/-^* mice, which also had a ≍15% reduction in flurothyl threshold relative to WT (Hawkins et al., 2021; Speca et al., 2014). In contrast, *Kcnb1^G379R^* mice with a dominant-negative variant had a more pronounced response to flurothyl compared to *Kcnb1^C/C^* mice. Compared to WT, *Kcnb1^G379R/G379R^* homozygotes had a 37% reduction in flurothyl threshold, while *Kcnb1^G379R/+^* heterozygotes had a ≍15% reduction (Hawkins et al., 2021). The magnitude of threshold reduction compared to these other *Kcnb1* lines suggests that *Kcnb1^C/C^*results in loss-of-function rather than a dominant-negative effect. Previous work showed that repetitive action potential firing was completely suppressed in cortical pyramidal neurons co-expressing WT and R306C channels (Saitsu et al., 2015). This, coupled with differences in mechanisms of seizure induction by KA versus flurothyl, may underlie the observed differential response in *Kcnb1^C/+^* heterozygotes. Flurothyl acts as a GABA antagonist, suppressing presynaptic GABA release, while KA acts on both pre-and post-synaptic receptors to suppress presynaptic GABA release *and* enhance postsynaptic activation of glutamatergic neurons (Falcón-Moya et al., 2018). Thus, despite the failure of action potential firing in heterozygous pyramidal neurons, the KA condition provides an excitatory drive that is absent in the flurothyl condition.

Although seizure susceptibility can give us a measure of altered excitatory/inhibitory balance, the role of KV2.1 as a dynamically regulated homeostatic modulator of neuronal excitability makes it challenging to directly translate these acute effects to epilepsy propensity. Thus, we performed EEG analysis to study epileptiform activity and background EEG properties. Among individuals with the *KCNB1*-p.R306C variant, common EEG findings are diffuse polyspike waves, high amplitude spike wave complexes and continuous spike and wave during sleep (CSWS) (de Kovel et al., 2017; Marini et al., 2017). *Kcnb1^C/+^* and *Kcnb1^C/C^* mice were found to have markedly abnormal ictal-interictal EEG patterns, including many of the above listed characteristics. Most evident in the EEG traces were lengthy, rhythmic, high-amplitude spike wave trains, sometimes lasting for hours, while the mouse was immobile. Although we cannot definitively conclude that the *Kcnb1^R306C^*model recapitulates CSWS, the similarity is striking. Previous work identified spike and wave complexes lasting up to 15 minutes in *Kcnb1^G379R^* mice, while spontaneous seizure activity was absent from recordings in *Kcnb1^-/-^* mice (Hawkins et al., 2021; Speca et al., 2014); thus, the frequency and duration of these complexes in *Kcnb1^R306C^* mice was unprecedented. The recently reported *KCNB1*-p.R312H mouse model also showed long duration seizure activity (4-5 min), defined by sustained rhythmic synchronous discharges, providing further support that the persistent spiking identified *Kcnb1^R306C^* is pathogenic (Bortolami et al., 2022). Future studies using high-dose benzodiazepines, valproate, or corticosteroids would be useful to determine if the rhythmic high-amplitude spike wave trains in the *Kcnb1*^R306C^ mouse model can be attenuated with drugs commonly used for CSWS (Sánchez Fernández et al., 2014).

Limitations of this study that were not in the scope of this manuscript and will be addressed in the future include the following. Our analysis of subcellular localization and AIS maturation was a snapshot at DIV16-18, and therefore do not preclude the possibility of impaired maturation occurring at a later time point. In addition, our EEG recordings did not include EMG or accelerometer data necessary for systematically staging sleep, an essential feature for CSWS diagnosis; thus, precluding definitive classification of the long bouts of rhythmic high-amplitude spike wave trains accompanied by behavioral immobility as CSWS. Finally, effects at the level of circuit, network, and non-conducting roles of KV2.1 were not probed in this study and future studies will be necessary to determine their contributions to clinical phenotypes.

## 5. Conclusions

In summary, we generated and characterized a novel knock-in mouse model of *KCNB1* encephalopathy due to voltage-sensor dysfunction of KV2.1 channels. This represents a major category of dysfunction observed for channelopathies and the model will be a valuable resource for understanding disease pathophysiology and evaluating potential therapies.

## Supporting information

List of Supplemental Material

## Abbreviations

AnkG: Ankyrin G
AIS: axon initial segment
flurothyl: Bis(2,2,2-trifluoroethyl) ether
CI: confidence interval
DIV: days in vitro
EEG: electroencephalogram
FFT: fast fourier transform
GTCS: generalized tonic-clonic seizure
gRNA: guide RNA
HDR: Homology directed repair
ICC: immunocytochemistry
IDT: Integrated DNA Technologies, Inc.
KA: kainic acid
LoF: loss-of-function
MJ: myoclonic jerk
PBS-T: PBS+tween-20
PAM: protospacer adjacent motif
RFLP: restriction fragment length polymorphism
RNP: ribonucleotide protein complex
ssODN: single-stranded donor oligonucleotide
SPF: specific pathogen free
WB: western blot
WT: wildtype

## CRediT authorship contribution statement

**Seok Kyu Kang:** Conceptualization, Methodology, Formal analysis, Investigation, Writing - Original Draft, Writing - Review & Editing, Visualization. **Nicole A. Hawkins:** Conceptualization, Methodology, Formal analysis, Investigation, Writing - Original Draft, Writing - Review & Editing, Visualization, Project administration. **Dennis-Echevarria-Cooper:** Investigation, Formal analysis, Writing - Review & Editing. **Erin M. Baker:** Investigation, Formal analysis, Writing - Review & Editing. **Conor J. Dixon:** Investigation, Formal analysis, Writing - Review & Editing. **Nathan Speakes:** Investigation, Formal analysis, Writing - Review & Editing. **Jennifer A. Kearney:** Conceptualization, Formal analysis, Investigation, Writing - Review & Editing, Visualization, Project administration, Funding acquisition.

## Declaration of Competing Interest

The authors declare no competing interests related to this study

## Data Availability

Data will be made available upon reasonable request.

## Acknowledgements

We thank *KCNB1* families for their support, and Colette Brill-Forman for technical assistance. The genetically engineered mice were generated with the assistance of Lynn Doglio and Eugene Wyatt in Northwestern University Transgenic and Targeted Mutagenesis Laboratory. Imaging work was performed at the Northwestern University Center for Advanced Microscopy generously supported by NCI CCSG P30 CA060553 awarded to the Robert H Lurie Comprehensive Cancer Center. This work was supported by generous philanthropic donations and the National Institutes of Health grant U54 NS108874 (JAK).

## Notes

### Competing Interest Statement

The authors have declared no competing interest.

