## Supplementary material for "Altered neurological and neurobehavioral phenotypes in a mouse model of the recurrent *KCNB1*-p.R306C voltage-sensor variant": List of Supplemental Material

for

### Altered neuronal and behavioral excitability in a mouse model of the recurrent *KCNB1*-p.R306C voltage-sensor variant

**List of Supplemental Materials**

1. **Supplemental Video S1**. Generalized tonic-clonic seizure in a *Kcnb1*$^{C/C}$ mouse. <Suppl Video S1 Kcnb1-R306C-Homozygote.mp4>

2. **Supplemental Video S2**. Video-EEG of a spontaneous generalized tonic-clonic seizure in a *Kcnb1*$^{C/+}$ mouse. This video corresponds to the EEG trace presented in Figure 5a. <Suppl Video S2 Kcnb1-R306C-Het vEEG.mp4>
